# Localization and function of multivesicular-bodies that release exosomes in islet cells: dysregulation during type-2 diabetes

**DOI:** 10.1101/2023.04.13.536686

**Authors:** Veerabhadraswamy Priyadarshini, Prajakta Belekar, Lakshmi Kothegala, Nikhil R. Gandasi

## Abstract

Type-2 diabetes (T2D) is characterized by high blood glucose due to compromised insulin secretion from pancreatic β-cells. β-cells primarily comprise insulin-secreting large-dense-core-vesicles/insulin-secretory-granules (ISGs) and also multivesicular-bodies (MVBs). MVBs are vesicles of endosomal origin containing intraluminal vesicles, which upon fusion with the plasma membrane, secrete exosomes. These play a significant role in the physiology and pathology of T2D via intercellular communication. The role of MVBs and their influence on ISGs of β-cells or their characterization is yet to be uncovered. In our study, we characterized the role of MVBs by comparing them to largely well-characterized ISGs in β-cells. We compared the density, localization, and exocytosis of MVBs with ISGs in β-cells. For this, we developed a novel probe where we exploit the efficiency of tetraspanins CD63 and CD151 to label the MVBs in β-cells. We showed that the β-cells have a significantly higher density of ISGs than MVBs. MVBs and ISGs are spatially localized apart within β-cells. The proteins that localize with MVBs are different from the ones that localize with ISGs. Exocytosis of ISGs occurs at the periphery of the β-cells and takes significantly lesser time when compared to exosome release, which is non-peripheral and takes a longer duration. Further, we also observed a significant reduction in the density of ISGs and MVBs in T2D patients’ islets compared to healthy controls. Studying the effect of MVBs on insulin secretion in physiological and T2D conditions has huge potential. This study provides a strong basis to open new avenues for such future studies.

## Introduction

Type-2 diabetes (T2D) is a major metabolic disorder with increasing incidence and prevalence worldwide to the epidemic level being the ninth major cause of death worldwide (1). International Diabetes Federation has estimated the global prevalence of diabetes to be 9.3% (463 million people) in 2019, and it is estimated to rise to 10.2% (578 million) by 2030 and 10.9% (700 million) by 2045. About 50.1% of people with diabetes worldwide are undiagnosed (2, 3).

Dysfunction in the secretion of islet hormones is one of the first signs of T2D. Within the islets, insulin-secreting - β, glucagon-secreting – α, and somatostatin-secreting - δ-cell’s viability and normal functioning are affected during human T2D (4–7). In pancreatic β-cells, insulin is stored in membrane-bound secretory granules called insulin-secretory-granules (ISGs) (8). Secretion of insulin into the bloodstream is essential for maintaining glucose homeostasis. Healthy β-cells secrete extracellular vesicles (EVs) limiting the formation of islet amyloids, thus helping their survival. During diabetes, limited secretion of EVs is observed, correlating with the formation of islet amyloids leading to β-cell death (9). β-cell-derived exosomes were found to improve glucose tolerance, increase insulin content in mice with abnormal glucose tolerance (10).

Exosomes (nanosized vesicles, 35nm to 150nm), a type of extracellular vesicles, are released from multivesicular-bodies (MVBs) (11–13). In endocytic pathways, early endosomes mature into late endosomes, and inward budding of limiting membrane of maturing /late endosomes form MVBs. MVBs contain intraluminal vesicles released as exosomes to the extracellular milieu upon fusion of MVBs with the plasma membrane (11, 14, 15). These exosomes carry cargo which includes DNA, proteins, lipids, mRNA, microRNA, and long noncoding RNA (lncRNA)(16–19). The cargo of the exosomes acts as messengers and are taken up by the cells in an autocrine and paracrine manner, therefore mediating cell-cell communication (15, 20–24).

Exosomes are the widely studied major EV type secreted from several tissue types, including the pancreas, adipose tissue, liver, immunocytes and skeletal muscles (25–31). In β-cell derived EVs, the cargo was found to be altered in the case of T2D, thus, modulating insulin signaling and influencing disease development (32). Increased miR-29 and decreased miR-26a in β-cell derived EVs lead to impaired insulin signaling in neighbouring cells and induce chronic low-grade inflammation via macrophages and monocytes in circulation (33). EVs with altered miRNA (reduced miR-26a and NCDase, increased miR-375-3p and miR-21-5p) induce apoptotic signals in recipient β-cells in a paracrine manner leading to cell death and dysfunction, causing diabetes-related pathological changes (34).

MVBs and ISGs are two different vesicle types of β-cells that fuse with the plasma membrane leading to exocytosis. ISG secretion and trafficking have been extensively studied using confocal fluorescent microscopy and total internal reflection fluorescent microscopy (TIRF) by utilizing Neuropeptide Y (NPY) as a large-dense-core-vesicle marker (35–38). NPY, on expression in the pancreatic β-cells, labels the insulin-secretory-granules. Such secretion and trafficking studies of MVBs were limited due to the limited availability of probes to label MVBs in live cells. Tetraspanin labels used to label MVBs in other cell types have opened new avenues to understand MVB trafficking and secretion (39–42). Tetraspanins like CD151, CD81, CD82 and CD9 are crucial for exosome biogenesis and cargo sorting into MVBs (43). These tetraspanins are enriched in intraluminal vesicles and exosomes (44). Their usage as biomarkers for exosomes in HeLa cells shows no changes in normal trafficking and secretion of MVBs (39, 42). The kinetics of release in non-secretory cell types including their arrival to membrane during docking has provided insights on their behaviour (41). The importance of these MVBs in pancreatic β-cells with regard to metabolic disorders has not been well studied. To understand this, we exploited CD63 for labelling MVBs in β-cells were explored to visualize trafficking and exocytosis of these vesicles. Further, we studied if MVB trafficking and fusion is similar to that of ISG by utilizing imaging techniques. We used a secretory cell type such as pancreatic β-cells, where correlation between ISGs and MVBs (source of exosomes) could be visualized. We were able to compare ISGs and MVBs with respect to their density, localization and exocytosis in physiological conditions and their density in case of T2D. Surprisingly, we saw that the density of both MVBs and ISGs was decreased in case of T2D, showing that insulin secretion and survivability of islet cells are compromised during T2D due to EV release. This study will open up avenues for a better understanding of how they impact each other’s function and their effect on maintaining glucose homeostasis.

## Results

### Density and localization of MVBs and ISGs

To study MVB and ISG populations in pancreatic β-cells, we explored the strategies implied in previous literature for labelling MVBs (39, 40, 45–47) and LDCVs (38, 48). There are no promising markers to label and visualize the exocytosis of MVBs in pancreatic β-cells. We exploited tetraspanins CD63 and CD151 to label MVBs in β-cells based on their ability to label MVBs in other cell types (39–41). When CD63-EGFP and CD151-mEmerald were overexpressed in β-cells, we saw punctate structures near the plasma membrane of β-cells (Figure 1A-B), similar to what had been observed in HeLa cells (39). Similarly, we observed NPY-mCherry labelled ISGs near the plasma membrane of β-cells (Figure 1C), similar to what has been described before (49). Fluorescent puncta observed in single cells using TIRF microscopy was quantified using an image J Plugin “find maxima” (Figure 1D) (details in the methods). The density of CD63-EGFP and CD151-mEmerald labelled MVBs and NPY-mCherry (LDCV marker) labelled ISGs were normalized to the area of the cell. The density of CD63-EGFP and CD151-mEmerald was 0.074 μm^-2^ and 0.079 μm^-2^ respectively. No significant difference was observed between the density of MVBs labelled using CD63-EGFP and CD151-mEmerald. The density of ISGs observed was 0.439 μm^-2^ and the β-cells were found to contain significantly higher density of ISGs than MVBs p<0.001 (Figure 1E).

**Figure 1.**
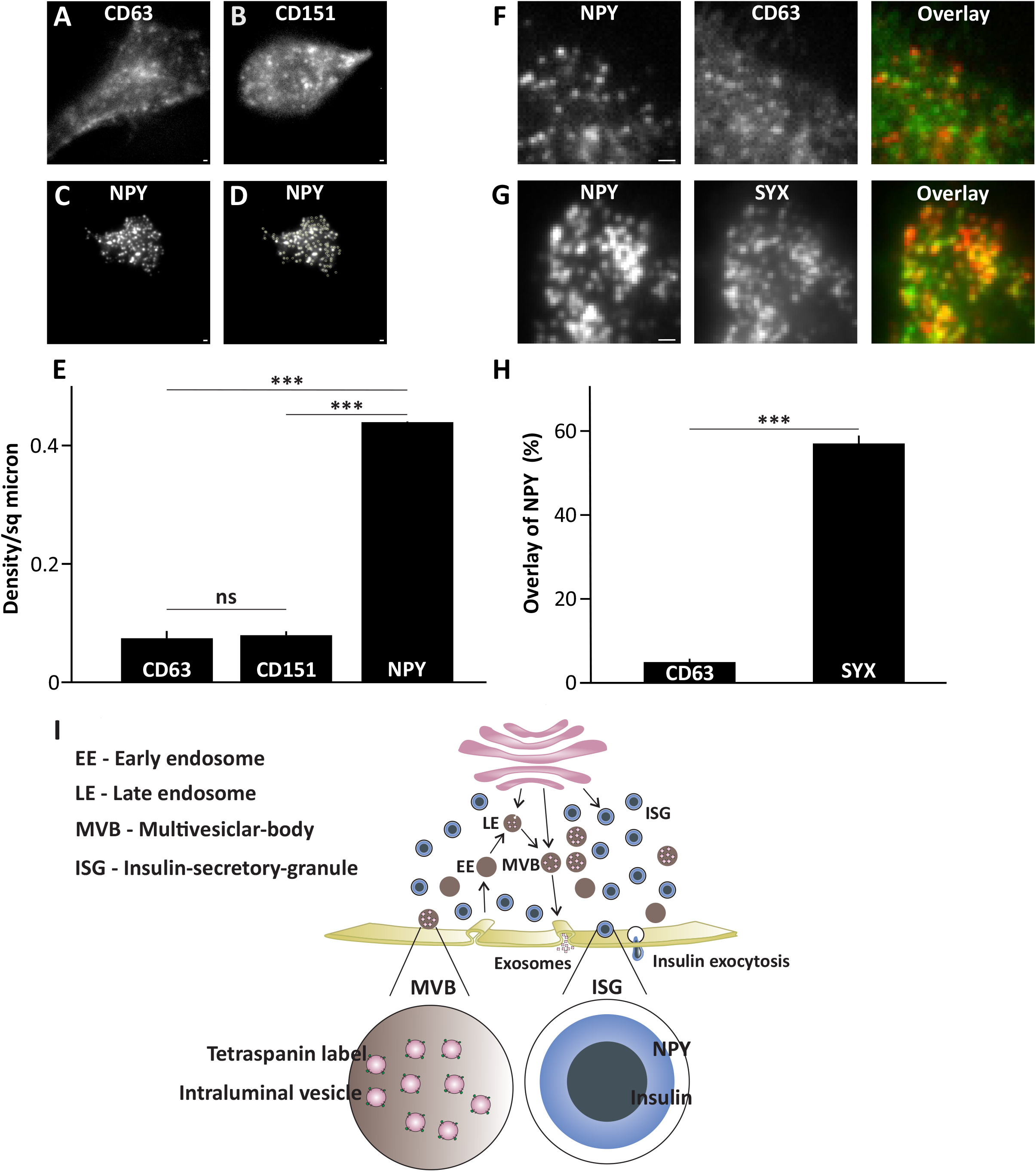
Density and localization of multivesicular-bodies (MVBs) and insulin-secretory-granules (ISGs). **A-B** Image of a cell expressing CD63-EGFP (A) or CD151- mEmerald (B) which labels MVBs. Scale bar 1μM. **C** Image of a cell expressing NPY-mCherry which labels ISGs. Scale bar 1μM. **D** Detection of ISGs based on the analysis described in methods. Scale bar 1μM. **E** Density of CD63-EGFP, CD151-mEmerald and NPY-mCherry (for images like in A-C). Data are presented as mean ± SEM for n = 19 cells (CD63-EGFP), n = 21 cells (CD151-mEmerald), n = 22 cells (NPY-mCherry) from at least 3 independent experiments in each case. ***p < 0.001 (details in the methods). **F** Images of cell expressing ISGs (NPY-mCherry) in red channel with MVBs (CD63-EGFP) in green channel with a corresponding image in the overlay. Scale bar 1μM. **G** Images of cell expressing ISGs (NPY-mCherry) in red channel with Syntaxin-1A-EGFP in green channels with a corresponding image in the overlay. Scale bar 1μM **H** Colocalization from images such as F-G calculated as described in the methods. Data are represented as mean ± SEM for n= 36 cells for each case from at least 3 independent experiments. ***p < 0.001. **I** Schematic representation of localization and trafficking of tetraspanin labelled MVBs and NPY labelled ISGs.

Further, we wanted to evaluate if the hotspots of exocytosis for ISGs are utilized for MVB exocytosis by analysing the localization of MVBs with ISGs near the plasma membrane. Cells co-transfected with NPY-mCherry and CD63-EGFP or NPY-mCherry and Syntaxin-1A-EGFP as control were imaged using TIRF microscopy. Syntaxin was used as a control since it is a t-SNARE, which facilitates the fusion of ISGs to the plasma membrane. Overlay of NPY-mCherry with CD63-EGFP (Figure 1F) or Syntaxin-1A-EGFP (Figure 1G) was analysed for colocalization using Metamorph (details in methods). The percentage colocalization of NPY-mCherry with CD63-EGFP was found to be 4.96% where as it was 57.03% for NPY-mCherry colocalizing with Syntaxin-1A-EGFP (Figure 1H). This data suggests that NPY-mCherry labelled ISGs are spatially apart from CD63-EGFP labelled MVBs. These were compared to positive control NPY-mCherry, which had a significantly high degree (p<0.001) of overlap with Syntaxin-1A-EGFP (Figure 1H). We conclude that the density of MVBs is significantly lesser than ISGs and they are spatially localized apart in β-cells. Therefore, they cannot utilize the exocytosis machinery of each other.

### Exocytosis of MVBs and ISGs have distinct dynamics

We visualized the exocytosis of MVBs and ISGs to compare the kinetics of fusion and release of both the vesicle types in β-cells. We imaged the cells transfected with CD63-EGFP or NPY-mCherry, to visualize the exocytosis of MVBs and ISGs, respectively, using TIRF microscopy. Cells were stimulated for MVB and ISG release (details in the methods), where individual fluorescent puncta disappear in one frame. Such events were identified as release events in the series of time-lapse images captured based on the criteria: a) disappearance of fluorescence in a single frame b) sudden loss of fluorescence without any reappearance in the same region during the time duration of the experiment. These criteria were followed for detection of MVB (CD63-EGFP) and ISG (NPY-mCherry) exocytosis (Figure 2A-B). When the single release event of MVBs (Figure 2C-D) and ISGs (Figure 2E) were plotted as ΔF (details in the methods) over time, we see that the release of MVBs is slower than that of ISGs (Figure 2C-E). We calculated the average release time by analysing multiple release events of MVBs and ISGs and plotted the same (Figure 2H). It was observed that average release time for MVB is above 40s and same is less than 10s for ISGs after stimulation. It is clear that MVBs take significantly higher time for release when compared to ISGs (p<0.001) (Figure 2H). We plotted a histogram based on the fusion events of MVBs (Figure 2F) and ISGs (Figure 2G) upon stimulation (stimulation starts 10s after beginning of the experiment, details in the methods), we see that most of the release events both in case of MVBs and ISGs start within 10s after stimulation (Figure 2F-G). The number of fusion events were significantly lower in the case of MVBs (0.017 ± 0.005), when compared to ISGs (0.058 ± 0.010). These data indicate that MVBs and ISGs follow distinct release kinetics.

**Figure 2.**
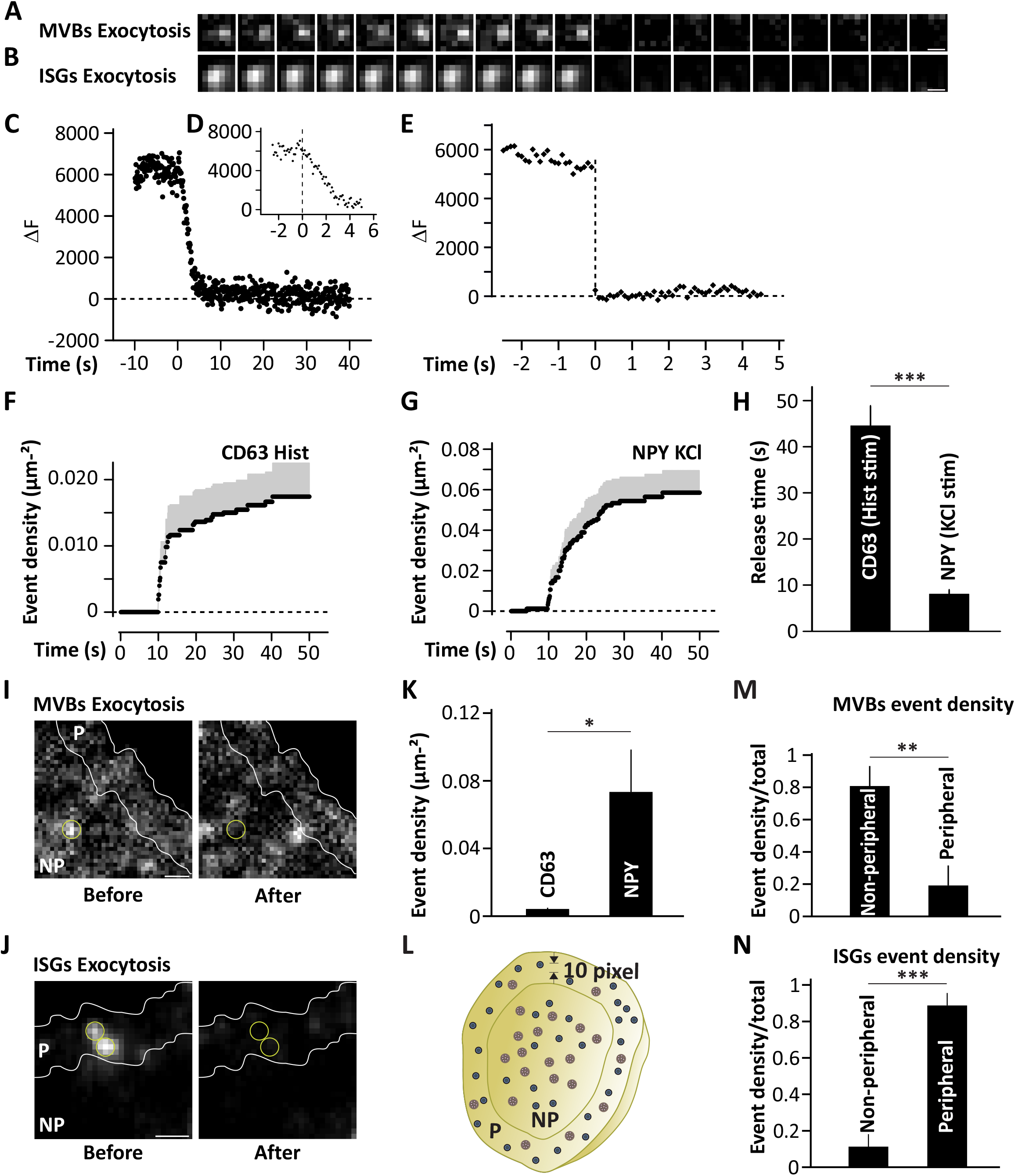
Exocytosis of MVBs and ISGs. **A** Time series strip showing disappearance of MVBs from the plasma membrane due to exocytosis. Scale bar 0.5μM. **B** Same as A for ISGs. Scale bar 0.5μM. **C** Fluorescence from single exocytotic event of MVBs over a period of 40s showing the kinetics of release. Fluorescence is quantified using ΔF as described in methods. **D** Same as C over a period of 6s. **E** Same as C for single exocytotic event of ISGs over a period of 5s. **F** Density of exocytosis events of MVBs after stimulation with histamine. The stimulation was initiated at 10s. **G** Same as F for exocytotic events of ISGs after stimulation with KCl. **H** Average release time for the exocytosis of MVBs and ISGs based on kinetics plotted for events similar to the ones plotted in D-E. Data are presented as mean ± SEM for n= 9 cells for each case. ***p < 0.001. **I** Image showing MVB exocytosis event, before (left) and after (right) loss of fluorescence of CD63-EGFP (MVBs) as peripheral (P, within 10 pixels from the border of the cell) and non-peripheral (NP, at least 10 pixels away from the border of the cell) events as described in methods. Scale bar 1μM. **J** Same as I for ISGs. Scale bar 1μM. **K** Density of total exocytosis events of MVBs and ISGs. data are represented as mean ± SEM for n= 8 cells for each case. *p < 0.05. **L** Schematic drawing showing peripheral and non-peripheral region of a cell considered for analysis. **M** Density of peripheral and non-peripheral events as fraction of total events observed during exocytosis of MVBs. **p < 0.01. **N** Same as M for ISGs. ***p < 0.001.

### MVBs and ISGs have preferential site of release

The observed difference in the localization and release kinetics of MVBs and ISGs led us to look for their release site within the cell. Our previous results described that MVBs and ISGs are spaced apart in non-stimulatory conditions (Figure 1H). We further evaluated if they remained apart during stimulated conditions as well. To evaluate the distribution and density of release events, we classified the labelled cell into peripheral (within 10 pixels from the border) and non-peripheral regions (after 10 pixels from border towards centre) (Figure 1L). We identified MVB and ISG release events based on the criteria (more details in the methods) mentioned above (Figure 2A-B).

The release events identified in peripheral and non-peripheral regions of a cell are shown as before and after exocytosis of MVBs (CD63-EGFP, Figure 2I) and ISGs (NPY-mCherry, Figure 2J), respectively. The total number of release events of MVBs and ISGs were normalized to the area of the cell to calculate the density of the distribution of events. We see that the density of release events overall for MVBs (0.004 μm^-2^) was significantly lower (p<0.05) when compared to the ISG event density (0.073 μm^-2^) (Figure 2K). We further evaluated how many of the total events are localized to peripheral or non-peripheral regions (as described previously) and expressed it as fraction of peripheral and non-peripheral events of MVBs and ISGs respectively. MVBs were released preferentially from the non-peripheral region (total events = 0.003 μm^-2^, fraction non-peripheral = 0.808/total) when compared to the peripheral region (total events = 0.001 μm^-2^, Fraction peripheral = 0.192/total) (p<0.01) (Figure 2M). On the other hand, ISGs released significantly (P<0.001) more from the periphery of the cells (0.005 μm^-2^, 0.1125/total) when compared to the non-peripheral release events (0.069 μm^-2^, 0.8875/total) (Figure 2N). Therefore, MVBs release preferentially from non-peripheral regions and ISGs from peripheral regions, indicating their distinct release site during exocytosis in β-cells.

### Endocytotic proteins are colocalizing with MVBs

To analyse the internalization of the exocytosis machinery soon after the MVB fusion and release of cargo, we looked for the association of a few endocytotic proteins. This gives an idea of how fast sustained release events can occur over a time period. Fluorescently tagged endocytotic proteins Clathrin and NECAP (adaptin ear-binding clathrin-associated protein) and endolysosomal protein LAMP1 (Lysosomal Associated Membrane Protein 1) were analysed for their localization with MVBs. Cells co-transfected with CD63-EGFP and NECAP-mCherry or CLC (clathrin light chain)-mCherry or LAMP1-mCherry were imaged using confocal microscopy. We observed CD63-EGFP puncta overlapping with fluorescent puncta of NECAP-mCherry (Figure 3A) or CLC-mCherry (Figure 3B) or LAMP1-mCherry (Figure 3C). Overlay of CD63-EGFP with NECAP-mCherry or CLC-mCherry or LAMP1-mCherry was analysed for colocalization (using Image J as described in methods). We see that CD63-EGFP colocalized with NECAP at 43.81 % and same was 40.14 % with CLC and 40.44% with LAMP1-mCherry (Figure 3D). The colocalization indicates that these endocytotic proteins do interact with CD63, suggesting the involvement of endocytotic proteins in the internalization of fusion machinery for quick recycling and sustained release.

**Figure 3.**
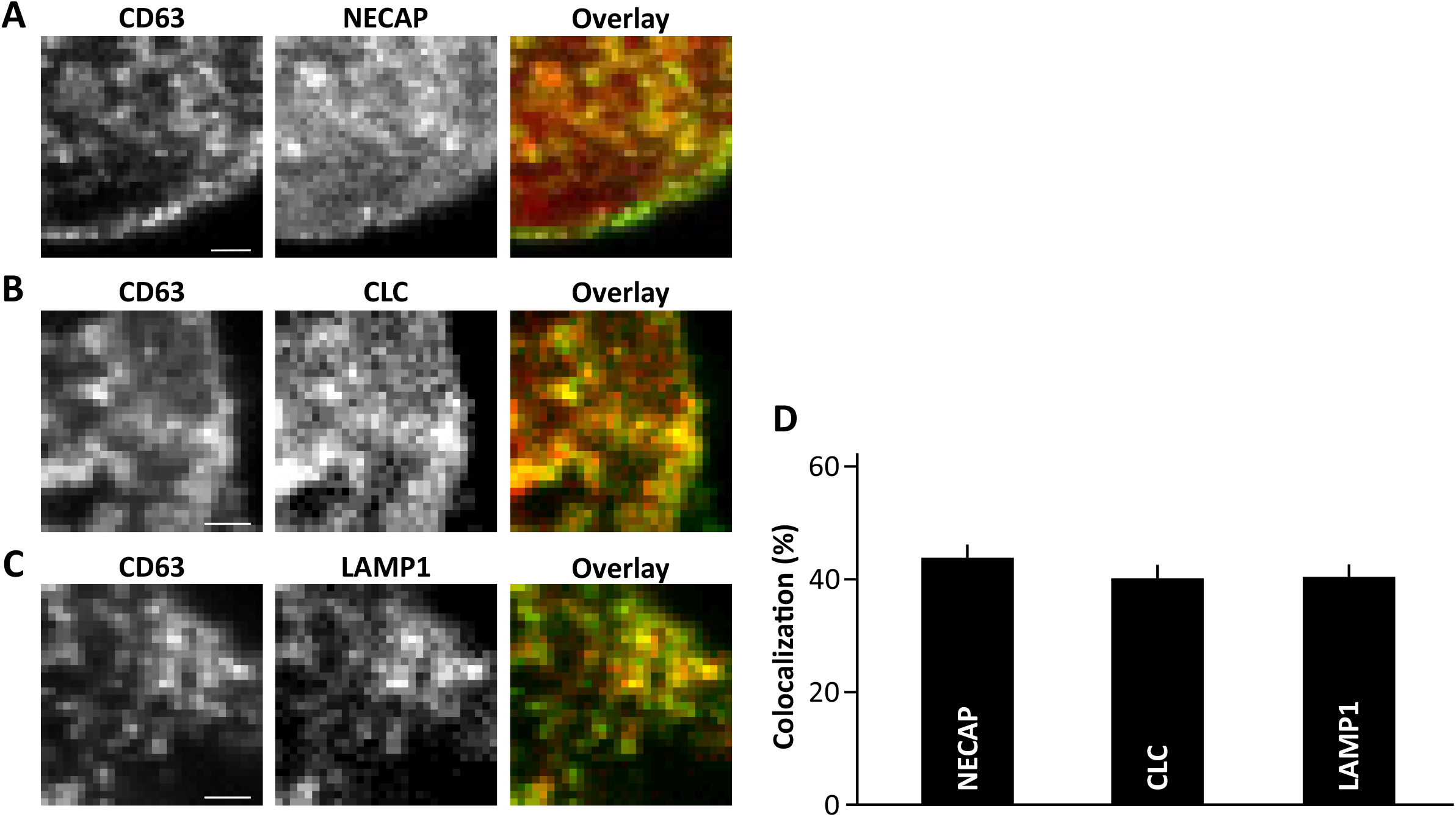
Colocalization of MVBs with endocytotic proteins. **A-C** Images of cell expressing MVBs (CD63-EGFP) in green channel with NECAP-mCherry (A), CLC-mCherry (B), or LAMP1- mCherry (C) in red channel with a corresponding image in the overlay. Scale bar 1μM. **D** Colocalization of MVBs (CD63-EGFP) with NECAP-mCherry, CLC-mCherry and LAMP1-mCherry measured as described in the methods. Data are presented as mean ± SEM for n = 14 cells (NECAP), n =17 cells (CLC), n =21 cells (LAMP1) from at least 3 independent experiments in each case.

### MVBs and ISGs are reduced in type-2 diabetes

Extrapolation of the density of vesicle types observed in β-cell line to the human pancreatic islets is more relevant to study the alterations during disorders such as T2D. The density of MVBs and ISGs in pancreatic islet β-cells of human donors with non-diabetic (ND) and T2D conditions were evaluated for this purpose. ND and T2D pancreatic islet β-cells were transduced with CD63-mCherry or NPY-mCherry to label MVBs or ISGs, respectively. We observed CD63-mCherry positive puncta labelling MVBs (Figure 4A-B) or NPY-mCherry positive puncta labelling ISGs (Figure 4D-E) in human islet cells. The fluorescent puncta observed in both cases were quantified using an image J plugin “find maxima” (details in the methods). The density of vesicles in each case was calculated by normalizing the count to area of that particular islet cell.

**Figure 4.**
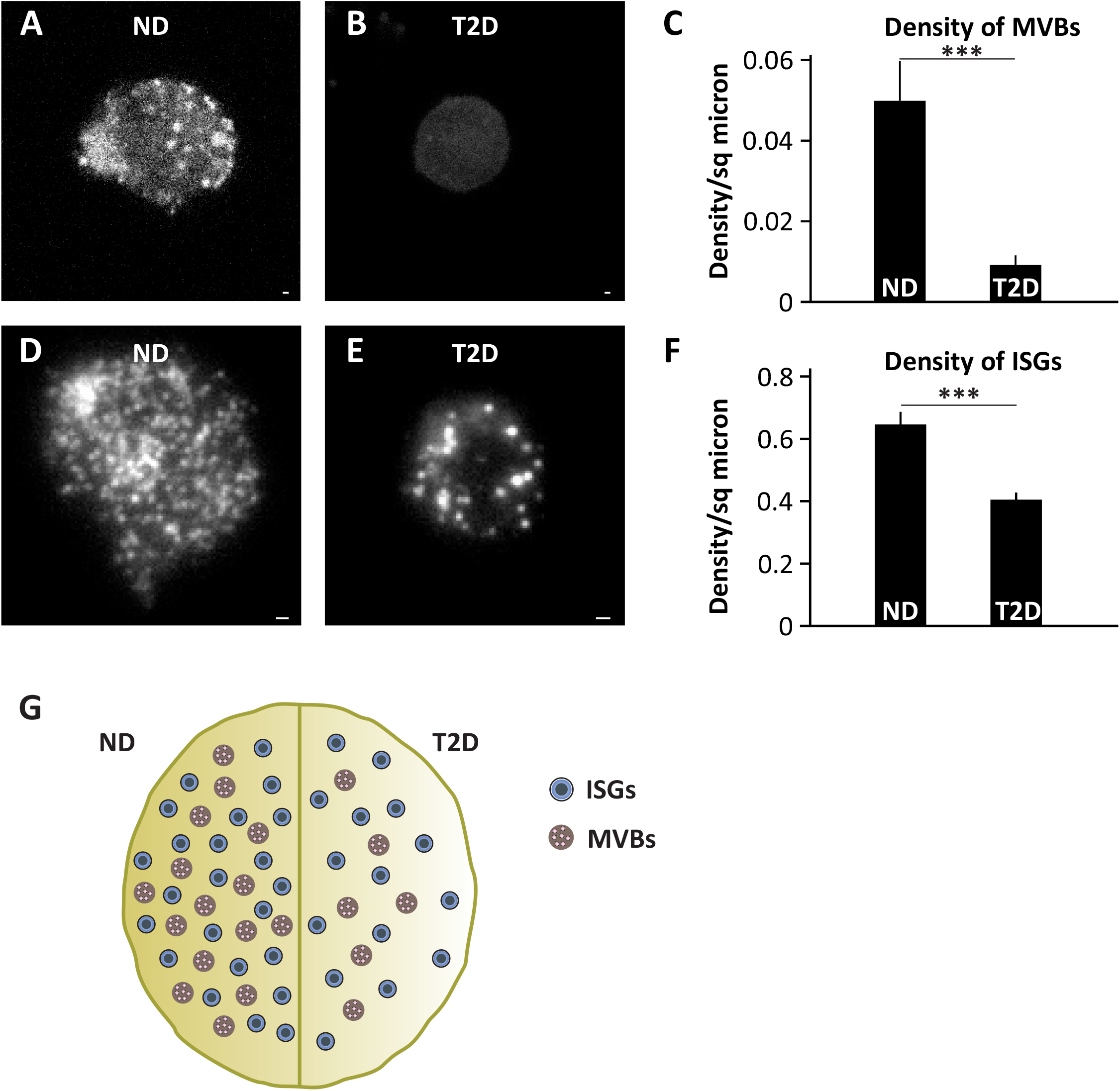
Comparison of MVBs and ISGS distribution in non-diabetic (ND) and type-2 diabetic (T2D) β-cells from human pancreas. **A-B** Image showing MVBs in ND (A) and T2D (B) human islet cells. Scale bar 1μM. **C** Density of MVBs in ND and T2D human islet cells. Data are presented as mean ± SEM. For n = 92 islets - T2D (3 donors), n = 21 islets - ND (2 donors), ***p < 0.001. **D-E** Same as A-B for ISGs. Scale bar 1μM. **F** Density of ISGs in ND and T2D human islets data are presented as mean ± SEM. For n = 84 islets - T2D (3 donors), n = 28 islets - ND (3 donors). ***p < 0.001. **G** Schematic diagram showing density of MVBs and ISGs in ND and T2D pancreatic islet cells.

In ND condition, we see a higher number of CD63-mCherry labelled MVBs (Figure 4A), but almost no MVBs in the case of T2D (Figure 4B). When quantified, a significant decrease in density of CD63-mCherry labelled MVBs was seen in T2D cells (0.009 μm^-2^) when compared to ND cells (0.049 μm^-2^) p<0.001 (Figure 4C). Similarly, NPY-mCherry labelled ISGs were observed in ND human islet cells (Figure 4D) and visually the number of these were decreased in case of T2D (Figure 4E). Quantifying the same in many such cells showed NPY-mCherry labelled ISGs were significantly decreased (p<0.001) in T2D (0.405 μm^-2^) when compared to ND condition (0.647 μm^-2^) (Figure 4F). The density of MVBs in ND human islet cells is lesser than ISGs, similar to our previous result shown in β-cell lines (Figure 1E). The decrease in density of MVBs and ISGs in diabetic human islet cells is maybe because of their non-functionality. This might reflect decreased synthesis and secretion from ISGs (35) or MVBs (9, 50) as shown previously.

## Discussion

Pancreatic β-cells contain large-dense-core-vesicles (ISG) that secrete insulin to maintain glucose homeostasis. The secretory granules are ∼300nm in diameter, and a single pancreatic β-cell consists of more than 1000 ISGs (51). NPY, a marker of large-dense-core-vesicles has been used here to label ISGs of pancreatic β-cells (49). Pancreatic β-cells synthesize and secrete exosomes, in addition to insulin secretion. Tetraspanins like CD63 and CD81 enriched on intraluminal vesicles or exosomes are well established as a marker of isolated exosomes in other cell types (43). It includes non-secretory cells like HeLa cells, where tetraspanins have been exploited to label MVBs containing intraluminal vesicles (ILVs) and visualized in real time (39). The same has not been characterized in pancreatic β-cells, a secretory cell type where the populations and function of both MVBs and secretory large-dense core vesicles such as ISGs might vary compared to a non-secretory cell. In this study, we have exploited tetraspanins like CD63 and CD151 as markers of MVBs and NPY as a marker of ISGs and observed that ISGs are highly populated vesicles in pancreatic β-cells when compared to MVBs. We also successfully employed CD63 to visualize the MVBs of pancreatic β-cells isolated from human islets. This will pave the way for understanding the exosome secretion in single human pancreatic β-cells and its relation to human diabetes.

Early endosomes that bud from plasma membrane either recycle back the proteins to the membrane or mature into late endosomes. These late endosomes acquire intraluminal vesicles (ILVs) to become MVBs (12, 52). MVBs can either fuse with plasma membrane to release exosomes, or mature into, or fuse with lysosomes (14). Tetraspanins like CD63, are enriched on ILVs of MVBs and also found on lysosomes. CD63, which is transported to the endosomes, will later make its way to MVBs and lysosomes (53). Since the life cycle of CD63 spans across various cellular organelles, we checked for the localization of CD63 with markers of endocytosis. Here, in pancreatic β-cells, CD63 is colocalizing with endocytotic proteins like clathrin and NECAP and a lysosomal protein LAMP1, which confirms its occurrence in biosynthetic and endolysosomal pathways. These data further confirm the origin and recycling of MVBs labelled with CD63.

On the other hand, ISGs are trafficked to the plasma membrane for fusion and exocytosis. Secretion of ISGs happens in a biphasic manner; the two phases of insulin secretion correspond to the exocytosis of different functional granule pools. A subset of granules, primed with fully assembled exocytotic machinery, constitutes for the readily releasable pool (8, 54), whereas, majority of the ISGs are localized deep inside the β-cells constituting the reserve pool. Exocytosis of granules from the reserve pool would require trafficking of these granules to the release site, docking of the vesicles to the plasma membrane, followed by priming (37, 49, 55) During SNARE-complex mediated exocytosis of ISGs, there is a fusion of VAMP2 (v-SNARE) with t-SNAREs’ Syntaxin-1A and SNAP-25 in the plasma membrane (37). The detailed mechanism of ISG secretion, exocytotic machinery, including accessory proteins involved, and the dynamics of ISG exocytosis is well studied and understood, but the same remains elusive in the case of MVB secretion. Trafficking and fusion of the MVBs to the plasma membrane is mediated by factors such as Rab GTPases and SNARE complexes (13, 52, 56). SNAREs like VAMP7 in the human leukemic cell line (K562), YKT6 in human embryonic kidney cells (HEK293) and human lung cancer cells (A549) were found to be the vesicular SNAREs involved in fusion. Target membrane SNAREs (t-SNAREs) like Syntaxin-1A in drosophila S2 cells and SNAP-23 in Hela cells were implied for fusion and secretion of MVBs (52, 56). The nature of the SNARE complex involved in the secretion is still not well understood. The detailed studies on the fusion machinery, including the accessory proteins, mechanism and dynamics of MVB secretion needs to be explored. There is high importance to exploring this in secretory cell types like β-cells. We chose to compare this with the secretion of ISGs and correlate it with the MVB secretion dynamics. There could be a high chance of having same exocytotic machinery for the fusion of both the vesicle types so that they are energetically favourable. Here, we assessed the possibility of MVBs and ISGs to use the same hotspots populated with proteins involved in the fusion machinery during exocytosis. On the contrary, MVBs and ISGs were localized apart near the plasma membrane; MVBs do not localize with syntaxin, which is a part of the fusion machinery for the release of ISGs (5.36% overlay of syntaxin with CD63). Therefore, fusion machinery of ISGs and MVBs are different, which leads us to evaluate the potential differences in the kinetics of exocytosis of these two vesicle types.

In this study, the tetraspanin CD63 established to label MVBs proved successful in analysing the kinetics of exosome exocytosis and the distribution of its release events in a single pancreatic β-cell in real-time (Figure 2). We were able to visualize the exocytosis of MVBs and compare it with the kinetics of ISGs marked with NPY in single β-cells. When we compared the exocytosis events of both vesicle types, we observed that MVBs take more time (>40s) while ISG exocytosis is rapid (<10s) (Figure 2H). Quantification of these events also revealed a lesser number of MVBs undergoing exocytosis than that of ISGs (Figure 2K). The difference observed was not only limited to its kinetics of release. We also noticed a difference in the preferential release site spatially arranged in single cells. Exocytosis of ISGs is generally localized to a small portion of pancreatic β-cells, referred to active region of release, suggesting the polarized arrangement of ISG exocytosis in β-cells, which we observe here as well (Figure 2J). These preferential release sites might aid in direct secretion into blood vessels (57). Although, ISG exocytosis is polarized (Figure 2J), MVBs are not the same. MVBs, on the contrary, release exosomes from non-peripheral regions (Figure 2I). Though ISGs and MVBs are being secreted from the same cell, the notable difference in their exocytosis is remarkable. Such differences in the exocytosis of these vesicle types could be because of different exocytosis machinery involved in their release.

Several studies have implicated the EV-mediated cross-talk affecting the β-cell function and/or viability (31, 58, 59). Healthy pancreatic β-cells and their normal functioning is essential for combating hyperglycaemia and insulin resistance seen during T2D development. β-cells secrete insulin via the exocytosis of ISGs to maintain the blood glucose, but the insulin secretion is affected during T2D, hence affecting the β-cell function and eventually viability. Islet cell function and viability are maintained due to the secretion of exosomes, which mediate cross-talk between other pancreatic β-cells and other islet cells (31). Exosomes secreted from healthy islets have a protective role in the survival and function of pancreas (58) by increasing insulin content during abnormal glucose tolerance (10). A subset of β-cells with high expression of CD63 shows enhanced insulin secretion. This subset was diminished in case of T2D (50). Similarly, islets from T2D patients had decreased secretion of EVs due to increased islet amyloids which could be reversed by adding EVs derived from healthy patients *in vitro* (9). These conditions trigger an immune response and hence alter the exosomal load in circulation (32). In agreement with the above studies, we observed a decreased density of MVBs and ISGs observed in the single islet β-cell of T2D patients. This shows that normal functioning at the single cell level leading to secretion EVs maintaining β-cell viability and insulin secretion is altered during T2D. These results show the potential of studying EVs for the diagnosis and treatment of T2D, while also providing insights on the synthesis and secretion of exosomes regulating cell to cell communication.

Uncovering the molecular mechanism behind exosome secretion has a unique edge in establishing MVB markers for studying the health of islet cells and understanding its secretion in relation to large dense core vesicle secretion, such as insulin. This will lay a foundation for using exosomes as diagnostic and prognostic tools to detect diabetes.

## Methods

### Cells

INS1 832/13 cells were cultured in RPMI 1640 (Invitrogen) supplemented with 10% fetal bovine serum (FBS), streptomycin (100 μg/ml), penicillin (100 μg/ml), sodium pyruvate (1 mM), and 2-mercaptoethanol (50 μM).

Transient transfections were performed on 25-mm poly-L-lysine–coated coverslips in 100 μL OptiMEM^®^ (Invitrogen) using 0.5 μL Lipofectamine® 2000 (Invitrogen), 0.2–0.6 μg plasmid DNA. The reaction was terminated after 3–5 h, and imaging was performed 24–48 h after transfection.

### Islet cell isolation and transduction

Human pancreatic tissue was obtained with the informed consent of the families involved, from the Nordic Network of Clinical Transplantation (Uppsala Regional Ethics Board ethical approval 2006/348) (35) or the ADI Isletcore at the University of Alberta (Alberta Human Research Ethics Board ethical approval Pro00001754) (60). Work involving human tissue complies with all applicable ethical standards for use in research, and the Gothenburg Regional Ethics Board, Sweden (098-18) and ethical committee at Indian Institute of Science, India (02/24.02.2023). The acquired tissue was cultured overnight and the islets were isolated (35). Trypsinization and plating of the isolated islets were performed to obtain single cells. The resulting single cells were maintained at 37 °C and 5% CO_2_ post being cultured in CMRL 1066 medium supplemented with 5.5 mM glucose, 10% FBS, 2 mM L-glutamine, and 1% penicillin-streptomycin. On the 22-mm poly-L-lysine-coated coverslips, cells were plated and given the night to settle. Adenovirus particles were added to the culture media at a concentration of 1:1000 and cultured for 24-36 hours before being imaged. The adenovirus particles were either adCD63-mCherry (developed by Genewiz, USA; details of the plasmid are provided in Verweij et al., 2018) or adNPY-mCherry (61).

### Solutions

Cells were imaged in a solution containing 138 mM sodium chloride (NaCl), 5.6 mM potassium chloride (KCl), 1.2 mM magnesium chloride (MgCl_2_), 2.6 mM calcium chloride (CaCl_2_), 3 mM D-glucose, 5 mM HEPES (pH 7.4 with 1M sodium hydroxide (NaOH).

For exocytosis of ISGs, the buffer instead contained 10 mM glucose and was supplemented with 2 mM forskolin and 200 μM diazoxide, a K+ ATP channel opener that prevents glucose-dependent depolarization. Exocytosis was then evoked by computer-timed local application of high K^+^ (75 mM KCl equimolarly replacing NaCl) through a pressurized glass electrode similar to those used for patch clamp experiments. Exocytosis of MVBs was evoked by the application of 100 μM histamine in a similar way as described for the high K^+^ above.

### Constructs

The constructs used in this study were CD63-pEGFP (obtained from addgene-62964), CD63-mCherry was created by replacing the pEGFP by using the multiple cloning sites in the CD63-pEGFP, CD151-mEmerald (obtained from addgene-54032), Neuropeptide Y (NPY)-mCherry (kindly provided by S Barg), Syntaxin-1A-EGFP (kindly provided by W Almers), NECAP-mCherry, CLC-mCherry and LAMP1-mCherry (kindly provided by late C Merrifield).

### Microscopy

Cells were imaged using a total internal reflection (TIRF) microscope based on an AxioObserver Z1 with a 100 ×/1.45 objective (Carl Zeiss, Jena, Germany). Excitation was from two DPSS lasers at 491 and 561 nm or individual lasers as described in individual experiments. The emission light was chromatically separated into separate areas of an EMCCD camera (Photometrics Evolve) using an image splitter (Photometrics DV2, Photometrics, Tucson, AZ, USA). Alignment of the two-colour channels was corrected as previously described (62).

The confocal microscope used for imaging was a Zeiss LSM780. The objective used was a 63x/1.40 (Zeiss). Excitation light of wavelength 561 nm (red, emission 578-696 nm) and 488 nm (green, emission 493-574 nm) was used. The size of the pinhole used was 0.61 μm, which corresponds to 1 Airy unit. Acquisition settings of imaging for the colours were set at 750 optical gain, 16-bit images with 0.11 μm/pixel resolution.

### Image Analysis

Density of the vesicles in INS1 832/13 cells, non-diabetic (ND) and type-2 diabetic (T2D) human islet cells were calculated using a script that used the built-in ‘find maxima’ function in ImageJ (http://rsbweb.nih.gov/ij) for spot detection (49). This count was then normalized to the area.

Visualization of exocytosis of vesicles in the form of their fluorescence changes was performed using the image analysis software MetaMorph (Molecular Devices, Sunnyvale, CA, USA). Fluorescence changes of exocytosis events were captured during the period of the time series. During exocytosis, there is a sudden disappearance of fluorescence within a few milliseconds as visualized in the TIRF field. These changes were treated with an algorithm implemented in the MetaMorph journal. This journal reads the average pixel fluorescence of the granule in 1) a central circle (c) of 3 pixels (0.5 μm) diameter, 2) a surrounding annulus (a) with an outer diameter of 5 pixel (0.8 μm). Granule fluorescence ΔF was obtained by subtracting the circle (c) with the annulus value (a) (ΔF=c-a). This value of ΔF was given as per-pixel average for the entire 0.5 μm-2 circle. ΔF valves of the vesicles was plotted against time (s) (38).

For identifying the exocytosis events as peripheral and non-peripheral, from the border of the cells, a region of 10 pixels were marked. Events observed as fluorescence change within the region of 10 pixels from the border were considered as peripheral events and the events in the region after 10 pixels from border towards the centre were considered non-peripheral events.

Colocalization was estimated using MetaMorph software. Regions of interest (ROIs) were marked manually in the red channel after identifying the vesicles. When these ROIs were transferred to the other channel, the centring was marked by a yes/no choice. This was based on the ROI positioning within one pixel at the centre of the previously identified ROIs. These were plotted as percentage colocalization after normalizing it to the area of the cell.

### Statistics

Data is presented as mean ± SEM unless otherwise stated. All the other data was tested for statistical significance using Students t-test for one-tailed, unpaired samples, as appropriate. Significant difference is indicated by asterisks (*p < 0.05, **p < 0.01, ***p < 0.001).

## Author Contributions

Conceptualization, N.R.G.; Methodology, N.R.G., L.K. and P.B.; Software, N.R.G.; Validation, L.K., and N.R.G.; Formal analysis, V.P., P.B. and N.R.G.; Investigation, V.P. and L.K.; Resources, L.K. and N.R.G.; Data curation, V.P., P.B. and N.R.G.; Writing—original draft, V.P. and N.R.G.; Writing—review & editing, V.P. and N.R.G.; Visualization, V.P., P.B., L.K. and N.R.G.; Supervision, L.K. and N.R.G.; Project administration, N.R.G.; Funding acquisition, N.R.G. All authors have read and agreed to the published version of the manuscript.

## Institutional Review Board Statement

Human islets were provided through the JDRF award 19-DSA-048 (ECIT Islet for Basic Research Program) and the Alberta Diabetes Institute Islet-Core, Canada and Nordic Network for Clinical Islet Transplantation (Uppsala), Sweden. Human islets are being utilized in the University of Gothenburg, Sweden and Indian Institute of Science, India as per ethics protocols numbered—098-18 from Regionala Etikprovningsnamnden Goteborg and 08/20 July 2022 from Institutional Human Ethics Committee (IHEC), Indian Institute of Science, India respectively.

## Acknowledgments

We would like to thank Dr. Saptadipa Paul for reading the manuscript and giving comments. We would also like to thank Chaitra N for helping us in data analysis and Eepsitha Marathe for helping with the cartoons. We thank Patrik Rorsman for his constant support and comments. We thank all the human donors who kindly provided their islets through the JDRF award 19-DSA-048 (ECIT Islet for Basic Research Program) and the Alberta Diabetes Institute Islet-Core and Nordic Network for Clinical Islet Transplantation (Uppsala).

## References

1. Zheng Y, Ley SH, Hu FB. Global aetiology and epidemiology of type 2 diabetes mellitus and its complications. Nat Rev Endocrinol. 2018;14(2):88–98.

2. Saeedi P, et al. Global and regional diabetes prevalence estimates for 2019 and projections for 2030 and 2045: Results from the International Diabetes Federation Diabetes Atlas, 9th edition. Diabetes Res Clin Pract. 2019;157:107843.

3. Makam AA, et al. Setting the Stage for Insulin Granule Dysfunction during Type-1-Diabetes: Is ER Stress the Culprit?. Biomedicines. 2022;10(11):2695.

4. Kothegala L, et al. Somatostatin Containing δ-Cell Number Is Reduced in Type-2 Diabetes. Int J Mol Sci. 2023;24(4):3449.

5. Fu J, et al. A glucose-dependent spatial patterning of exocytosis in human β-cells is disrupted in type 2 diabetes. JCI Insight. 2019;5(12):e127896.

6. Omar-Hmeadi M, et al. Paracrine control of α-cell glucagon exocytosis is compromised in human type-2 diabetes. Nat Commun. 2020;11(1):1896.

7. Thurmond DC, Gaisano HY. Recent Insights into Beta-cell Exocytosis in Type 2 Diabetes. J Mol Biol. 2020;432(5):1310–1325.

8. Rorsman P, Renström E. Insulin granule dynamics in pancreatic beta cells. Diabetologia. 2003;46(8):1029–1045.

9. Ribeiro D, et al. Extracellular vesicles from human pancreatic islets suppress human islet amyloid polypeptide amyloid formation. Proc Natl Acad Sci U S A. 2017;114(42):11127–11132.

10. Sun Y, et al. Exosomes from β-cells alleviated hyperglycemia and enhanced angiogenesis in islets of streptozotocin-induced diabetic mice. Diabetes Metab Syndr Obes. 2019;12:2053–2064.

11. Stoorvogel W, et al. The Biogenesis and Functions of Exosomes. Traffic 2002;3:321–330.

12. Raposo G, Stoorvogel W. Extracellular vesicles: Exosomes, microvesicles, and friends. Journal of Cell Biology 2013;200(4):373–383.

13. Kowal J, Tkach M, Théry C. Biogenesis and secretion of exosomes. Curr Opin Cell Biol. 2014;29(1):116–125.

14. Kalluri R, LeBleu VS. The biology, function, and biomedical applications of exosomes. Science. 2020;367(6478):eaau6977.

15. Simons M, Raposo G. Exosomes--vesicular carriers for intercellular communication. Curr Opin Cell Biol. 2009;21(4):575–581.

16. Colombo M, Raposo G, Théry C. Biogenesis, secretion, and intercellular interactions of exosomes and other extracellular vesicles. Annu Rev Cell Dev Biol. 2014;30:255–289.

17. Abels ER, Breakefield XO. Introduction to Extracellular Vesicles: Biogenesis, RNA Cargo Selection, Content, Release, and Uptake. Cell Mol Neurobiol. 2016;36(3):301–312.

18. Van Niel G, D’Angelo G, Raposo G. Shedding light on the cell biology of extracellular vesicles. Nat Rev Mol Cell Biol. 2018;19(4):213–228.

19. Zhang J, et al. Exosome and exosomal microRNA: Trafficking, sorting, and function. Genomics Proteomics Bioinformatics. 2015;13(1):17–24.

20. Huang-Doran I, Zhang CY, Vidal-Puig A. Extracellular Vesicles: Novel Mediators of Cell Communication In Metabolic Disease. Trends Endocrinol Metab. 2017;28(1):3–18.

21. Meldolesi J. Exosomes and Ectosomes in Intercellular Communication. Curr Biol. 2018;28(8):R435–R444.

22. Mathivanan S, Ji H, Simpson RJ. Exosomes: extracellular organelles important in intercellular communication. J Proteomics. 2010;73(10):1907–1920.

23. Février B, Raposo G. Exosomes: endosomal-derived vesicles shipping extracellular messages. Curr Opin Cell Biol. 2004;16(4):415–421.

24. Tkach M, Théry C. Communication by Extracellular Vesicles: Where We Are and Where We Need to Go. Cell. 2016;164(6):1226–1232.

25. Garcia-Martin R, et al. Tissue differences in the exosomal/small extracellular vesicle proteome and their potential as indicators of altered tissue metabolism. Cell Rep. 2022;38(3):110277.

26. Mei RY, et al. Role of Adipose Tissue Derived Exosomes in Metabolic Disease. Front Endocrinol (Lausanne). 2022;13:873865.

27. Mytidou C, et al. Muscle-derived exosomes encapsulate myomiRs and are involved in local skeletal muscle tissue communication. FASEB Journal. 2021;35(2):e21279.

28. Flaherty SE, et al. A lipase-independent pathway of lipid release and immune modulation by adipocytes. Science. 2019;363(6430):989–993.

29. Théry C, Ostrowski M, Segura E. Membrane vesicles as conveyors of immune responses. Nat Rev Immunol. 2009;9(8):581–593.

30. Sato K, et al. Exosomes in liver pathology. J Hepatol. 2016;65(1):213–221.

31. Chidester S, et al. The Role of Extracellular Vesicles in β-Cell Function and Viability: A Scoping Review. Front Endocrinol (Lausanne) 2020;11:375.

32. Freeman DW, et al. Altered extracellular vesicle concentration, cargo, and function in diabetes. Diabetes. 2018;67(11):2377–2388.

33. Sun Y, et al Expression of miRNA-29 in Pancreatic β Cells Promotes Inflammation and Diabetes via TRAF3. Cell Rep. 2021;34(1):108576.

34. Liu J, et al. Integrative biology of extracellular vesicles in diabetes mellitus and diabetic complications. Theranostics. 2022;12(3):1342–1372.

35. Gandasi NR, et al. Glucose-Dependent Granule Docking Limits Insulin Secretion and Is Decreased in Human Type 2 Diabetes. Cell Metab. 2018;27(2):470–478.e4.

36. Gandasi NR, et al. Ca2+ channel clustering with insulin-containing granules is disturbed in type 2 diabetes. J Clin Invest. 2017;127(6):2353–2364.

37. Omar-Hmeadi M, Idevall-Hagren O. Insulin granule biogenesis and exocytosis. Cell Mol Life Sci. 2021;78(5):1957–1970.

38. Barg S, et al. Syntaxin clusters assemble reversibly at sites of secretory granules in live cells. Proc Natl Acad Sci U S A. 2010;107(48):20804–20809.

39. Verweij FJ, et al. Quantifying exosome secretion from single cells reveals a modulatory role for GPCR signaling. J Cell Biol. 2018;217(3):1129–1142.

40. Mathieu M, et al. Specificities of exosome versus small ectosome secretion revealed by live intracellular tracking of CD63 and CD9. Nat Commun. 2021;12(1):4389.

41. Mahmood A, et al. Exosome secretion kinetics are controlled by temperature. Biophys J. 2023;S0006-3495(23)00131-5.

42. Sung BH, et al. A live cell reporter of exosome secretion and uptake reveals pathfinding behavior of migrating cells. Nat Commun. 2020;11(1):2092.

43. Andreu Z, Yáñez-Mó M. Tetraspanins in extracellular vesicle formation and function. Front Immunol. 2014;5:442.

44. Escola J-M, et al. Selective enrichment of tetraspan proteins on the internal vesicles of multivesicular endosomes and on exosomes secreted by human B-lymphocytes. J Biol Chem. 1998;273(32):20121–20127.

45. Verweij FJ, et al. The power of imaging to understand extracellular vesicle biology in vivo. Nat Methods. 2021;18(9):1013–1026.

46. Bebelman MP, et al. Real-time imaging of multivesicular body-plasma membrane fusion to quantify exosome release from single cells. Nat Protoc. 2020;15(1):102–121.

47. Verweij FJ, Revenu C, Arras G, et al. Live Tracking of Inter-organ Communication by Endogenous Exosomes In Vivo. Dev Cell. 2019;48(4):573–589.e4.

48. Gandasi NR, et al. Survey of red fluorescence proteins as markers for secretory granule exocytosis. PLoS One. 2015;10(6):e0127801.

49. Gandasi NR, Barg S. Contact-induced clustering of syntaxin and munc18 docks secretory granules at the exocytosis site. Nat Commun. 2014;5:3914.

50. Rubio-Navarro A, Gómez-Banoy N, Stoll L, et al. A beta cell subset with enhanced insulin secretion and glucose metabolism is reduced in type 2 diabetes. Nat Cell Biol. 2023;10.1038/s41556-023-01103-1.

51. Dean PM. Ultrastructural morphometry of the pancreatic -cell. Diabetologia. 1973;9(2):115–119.

52. Hessvik NP, Llorente A. Current knowledge on exosome biogenesis and release. Cell Mol Life Sci. 2018;75(2):193–208.

53. Pols MS, Klumperman J. Trafficking and function of the tetraspanin CD63. Exp Cell Res. 2009;315(9):1584–1592.

54. Bratanova-Tochkova TK, et al. Triggering and augmentation mechanisms, granule pools, and biphasic insulin secretion. Diabetes. 2002;51 Suppl 1:S83–S90.

55. Olofsson CS, et al. Fast insulin secretion reflects exocytosis of docked granules in mouse pancreatic B-cells. Pflugers Arch. 2002;444(1–2):43–51.

56. Xu M, et al. The biogenesis and secretion of exosomes and multivesicular bodies (MVBs): Intercellular shuttles and implications in human diseases. Genes Dis. 2022;

57. Qian WJ, et al. Detection of secretion from single pancreatic β-cells using extracellular fluorogenic reactions and confocal fluorescence microscopy. Anal Chem. 2000;72(4):711–717.

58. Xiao Y, et al. Extracellular vesicles in type 2 diabetes mellitus: key roles in pathogenesis, complications, and therapy. J Extracell Vesicles. 2019;8(1).

59. Salomon C, Das S, Erdbrügger U, et al. Extracellular Vesicles and Their Emerging Roles as Cellular Messengers in Endocrinology: An Endocrine Society Scientific Statement. Endocr Rev. 2022;43(3):441–468.

60. Lyon J, et al. Research-focused isolation of human islets from donors with and without diabetes at the Alberta Diabetes Institute IsletCore. Endocrinology 2016;157(2):560–569.

61. Meur G, et al. Insulin gene mutations resulting in early-onset diabetes: Marked differences in clinical presentation, metabolic status, and pathogenic effect through endoplasmic reticulum retention. Diabetes. 2010;59(3):653–661.

62. Taraska JW, et al. Secretory granules are recaptured largely intact after stimulated exocytosis in cultured endocrine cells. Proc Natl Acad Sci U S A. 2003;100(4):2070–2075.

